# Sexual selection leads to positive allometry but not sexual dimorphism in the expression of horn shape in the blue wildebeest, *Connochaetes taurinus*

**DOI:** 10.1101/2022.06.01.494318

**Authors:** Chlöe Gerstenhaber, Andrew Knapp

## Abstract

Sexual selection is thought to be an important driver of adaptation, speciation and extinction. Empirically testing these predictions across macroevolutionary timescales first requires an understanding of the morphology of secondary sexual traits in extant taxa. We used three-dimensional geometric morphometrics to analyse a large sample of the skull of the blue wildebeest, *Connochaetes taurinus*, in which horns are found in both sexes but only used in intrasexual competition in males. We show that the horns fit several predictions of secondary sexual traits; overall skull shape is significantly correlated with size (R^2^ = 0.38, *p* = 0.001), and the sexually selected horns show drastically higher growth rates and variation than any other skull element, supporting previous findings. We also find that despite showing significant sexual dimorphism in shape and size (R^2^ = 0.21, *p* = 0.001), allometric growth trajectories of sexes are identical (R^2^ = 0.01, *p* = 0.635) and dimorphism is not readily detectable without prior knowledge of sex, and is not possible when shape is corrected for size. Our results show that even with strong sexual selection operating in only one sex, the expression of secondary sexual traits may show characteristic and indistinguishable patterns of growth and variance in both sexes.

## Introduction

Sexual selection arises from competition for fertilisation opportunities and is responsible for the evolution of diverse ‘secondary sexual’ traits in the animal kingdom, including exaggerated morphologies, behaviours and strategies (Darwin, 1871; Andersson, 1994). Sexual selection is expected to have a powerful effect on evolution, speciation and extinction (Ritchie, 2007; Martínez-Ruiz and Knell 2016; Janicke et al., 2018), and although much theoretical and laboratory work has been done to test these predictions, exploring these effects over macroevolutionary timescales is more challenging. Incorporating fossil data into these studies is a possible solution but requires identification of sexually selected traits in fossil taxa; our incomplete knowledge of these taxa makes this difficult in practice (Knell et al., 2012). The biology of extant organisms is generally much better understood and can be used to explore the effects of sexual selection on morphology, which can then be extended to extinct datasets. An extant dataset can thus be a useful proxy for a fossil dataset while accounting for variables generally unknowable in fossil taxa, e.g. sex.

Secondary sexual traits are known to display characteristic patterns of growth and variation that distinguish them from functionally constrained naturally selected traits (Losos, 2011; Knell et al., 2012). For example, traits which act as visual signals often show positive static allometry, being proportionally larger in sexually mature individuals. This is revealed as a slope of greater than 1 when trait size is regressed against body size (O’Brien et al., 2018). This phenomenon is widely observed across the animal kingdom and is thought to arise by being an ‘honest signal’, because growing and maintaining proportionally larger traits is more efficient for larger individuals (Rodríguez and Eberhard, 2019; Somjee, 2021). Studies of secondary sexual traits in extant taxa tend to focus on the competing sex, which may impede their application to extinct taxa, where sex is often unknowable (Mallon, 2017; O’Brien et al., 2018; Cooper et al., 2019). Moreover, striking sexual dimorphism seen in many secondary sexual traits has led some researchers to suggest that such traits can only be identified by the presence of sexual dimorphism (Knell and Sampson, 2011; Borkovic and Russell, 2014). Nonetheless, sexual dimorphism in fossil taxa is difficult to detect and is hampered by our incomplete knowledge of extinct taxa (Mallon, 2017). Furthermore, examples exist of extant taxa in which a secondary sexual trait is expressed in both sexes of a species but may only perform a sexually selected role in one, further complicating the identification of these traits (Stankowich and Caro, 2009; Tobias et al., 2012).

Differing roles of secondary sexual traits between sexes may lead to selection for positive allometry in these traits being relaxed in one sex, leading to dimorphism in growth and variation between sexes (Evans et al., 2018; Tidière et al., 2020). Conversely, some secondary sexual traits may also perform a range of functions connected with social selection, which may operate in a similar way to sexual selection in both sexes (West-Eberhard, 1979; Tobias et al., 2012). Furthermore, there is a tendency in studies of sexually selected traits to focus on a single ‘focal’ trait, while other aspects of anatomy are neglected, which may bias results towards such ‘exaggerated’ traits (Bonduriansky, 2007). Traditional one-dimensional linear measurements of trait size are often employed in studies of sexually selected traits, but univariate data cannot account for more complex changes in shape with size or in differences in growth elsewhere in the organism (Goswami et al., 2019). The exaggerated growth and high variation typical of secondary sexual traits (O’Brien et al., 2018) suggest that they are largely free of the functional constraints of other, naturally selected traits, and consequently they are likely to be weakly integrated with the rest of the organism (Klingenberg, 2008). Phenotypic modularity, the tendency of sets of traits to form integrated ‘modules’ which covary more strongly than with other integrated sets of traits, can further help to determine interactions between traits by assessing their integration across an organism (Zelditch and Goswami, 2021). Sexually selected traits, such as horns, should form distinct modules because relaxed integration with the rest of the skull would allow them to respond to selection with some degree of independence (Klingenberg, 2008). Determining integration of traits across an organism may therefore be an important indicator of function, especially when attempting to identify sexually selected traits. Modern geometric morphometric (GM) techniques allow the analysis of shape across a number of associated traits and can therefore avoid bias towards a single trait (Goswami et al., 2019).

The ruminant family Bovidae comprises 143 extant species. The males of all species bear sexually selected horns which are constructed of a bony core, the os cornu, covered by a horn sheath formed of keratinised epidermis (Bubinek and Bubinek, 1990). Horn shape appears to be correlated with male fighting style in intrasexual competition (Caro et al., 2003). Although female bovids do not physically compete for mates and are therefore not expected to be subject to sexual selection in the same way as males, female horns are found in roughly half of all extant bovid species (Packer, 1983). It is thought that female bovid horns are maintained by either natural selection (e.g. predator defence), social selection (e.g. territoriality, male mimicry), or by genetic linkage to male horns (Stankowich and Caro, 2009). Many bovid taxa are well understood and specimens are readily available in museum collections, making them an ideal study group for investigating intraspecific variation of secondary sexual traits. The blue wildebeest, *Connochaetes taurinus*, is a medium-sized bovid in the tribe Alcelaphini (Castelló, 2016). Sexes resemble each other, with females being generally smaller than males and with less robust horns (Estes 2014, Castelló, 2016, Tidière et al., 2017). *C. taurinus* is divided into five subspecies spread across East and Southern Africa, often living in very large populations which can in turn lead to intense competition between males for mates. Male *C. taurinus* hold small territories which they defend with ritualised aggressive behaviour, and horns are used in these aggressive displays and in physical competition with other males, with larger males generally being dominant (Estes, 2014). The prominent role of horns in these contests suggest that, although obvious weapons, they may also have an important display function and thus may display the predicted positive allometry and increased variance of a secondary sexual display trait (O’Brien et al., 2018; Rodríguez and Eberhard, 2019). There is some disagreement over the role of horns in female *C. taurinus*, with predator defence and male mimicry being leading hypotheses, and they are not known to be used in physical competition between females (Estes, 2014). Although the horns of *C. taurinus* are known to show positive allometry in length in males (O’Brien et al., 2018), it is not known if this relationship is seen in females, nor how shape is related to size either in horns or in the skull in general.

Using three-dimensional (3D) geometric morphometrics, we analyse a large sample of *C. taurinus* skulls of both sexes and a range of sizes to test the following predictions:

1. Skull shape in *C. taurinus*, including horns, is significantly different between sexes

2. The horns of *C. taurinus* form a distinct phenotypic module which is weakly integrated with the rest of the skull

3. The horns are positively allometric with skull size, and at a higher degree than any other skull element

4. Allometric trajectories of males and females are significantly different

Gaining a thorough understanding of the patterns of growth and shape variation of the horns of *C. taurinus*, how the horns integrate with the rest of the skull, and how secondary sexual traits differ between sexes, will resolve questions which are often overlooked or poorly understood in studies of sexually selected traits. Ultimately, the answers to these questions can be used to better understand how to detect signals of sexual selection in the morphology of both extant and extinct taxa.

## Methods

A total of 75 *C. taurinus* skulls (Supplementary table S1) were digitised from the collections of the Natural History Museum, London (NHM, n = 73) and the Museum Für Naturkunde, Berlin (MfN, n = 2) using photogrammetry (Agisoft Photoscan, v. 1.4.3). Meshes were decimated to one million faces and landmarks were placed on the right half of each mesh using Stratovan Checkpoint (2020, v. 2020.10.13.0859). A total of 49 anatomical landmarks and 50 semilandmark curves were used to capture shape across the right side of the skull. Landmarks were placed on the keratinous horn sheath because it is present in all specimens and not removable. Additional surface semilandmarks were placed on a template specimen and projected to all other specimens in the R (v4.1.2; R core team, 2021) package *Morpho* (Schlager, 2017) to capture surface shape variation across the skull and horns (Bardua et al., 2019b), giving a total of 849 fixed and semilandmarks. Landmarks were estimated using the thin plate spline (TPS) method in the R package *Morpho* (Schlager, 2017) in specimens where minor damage prevented the placing of some landmarks. Several specimens (n = 5) were more severely damaged or incomplete and were thus omitted from the surface semilandmark dataset. Semilandmarks were slid to minimise bending energy. Analyses were performed on all surface semilandmarked specimens (n = 70) unless otherwise stated. All additional analyses performed on subsets of the dataset are presented in the supplementary data. Landmarks were reflected across the sagittal plane to reduce lateral inaccuracies in aligning the bilaterally symmetrical skull (Knapp et al., 2021). Specimens were then aligned using a generalised Procrustes alignment (GPA) in the R package *geomorph* (Adams and Otárola-Castillo, 2013), and the reflected landmarks were removed, leaving the original, Procrustes-aligned right-side landmarks for analysis. All further analyses were performed using R Statistical Software (v4.1.2; R Core Team 2021).

A principal components analysis (PCA) was performed on the Procrustes-aligned dataset to determine major trends in shape variation. Sexual shape dimorphism was assessed with a multivariate analysis of variance (MANOVA), using known sex of each specimen as the independent grouping factor. Two approaches were used to determine clustering in the shape data; one to assess grouping accuracy of sexes using *a priori* knowledge of specimen sex, and a second to assess number of clusters in the data. Firstly, a k-means cluster analysis (Hartigan and Wong, 1979) was performed on the Procrustes-aligned shape data, with the *k* value (i.e. number of expected groups) set at 2, representing two expected sexes. The k-means cluster analysis was repeated on centroid size (defined as the sum of the squared distance of each landmark to the geometric centre of each specimen, Zelditch et al., 2004)for all specimens, with *k* value again set at 2. Secondly, the optimum number of clusters in the shape data was assessed with two methods using a k-means approach without prior assumptions of group number, the average silhouette method (Rousseeuw, 1987) and the gap statistic method (Tibshirani et al., 2001). The entire dataset and subsets of the data containing each of the sexes alone were analysed with these approaches to determine two-group and no-group datasets could be distinguished. Finally, Hartigans’ dip test was performed on the first 8 principal components of the Procruste-aligned shape data to test for non-unimodality with the R package *diptest* (Maechler, 2016). To test for non-unimodality in specimen size, the dip test was also performed on the centroid sizes of all specimens.

Phenotypic modularity was assessed by comparing seven modularity hypotheses, comprised of subsets of the landmark data (supplementary data). Two methods were used; the maximum likelihood (ML) approach implemented in the R package *EMMLi* (Goswami and Finarelli, 2016), and by comparing the covariance ratios of the module hypotheses with the compare.CR function in *geomorph* (Adams and Otárola-Castillo, 2013). The globally-aligned modules identified in these analyses were subjected to a k-means cluster analysis to assess grouping by sex for each module.

Allometry in the dataset was explored by regressing shape with size, using the function procd.lm in the R package *geomorph* (Adams and Otárola-Castillo, 2013). Differences in allometric slope between sexes was assessed by including sex as a factor in the regression. Allometry analyses were repeated for each globally-aligned module defined in the modularity analysis. Additionally, centroid sizes of each module were regressed against whole-skull centroid size to assess size-based allometry, comparable to the more traditional approach employed by O’Brien et al. (2018). Shape data were corrected for allometry by removing the shape regression score calculated during the allometry regression to produce a size-independent shape dataset (Mitteroecker et al., 2004). Sexual dimorphism was assessed in the allometry-corrected dataset by repeating the dimorphism analyses performed on the raw shape data. Specifically, a MANOVA was performed using sex as the independent grouping factor, a k-means cluster analysis was performed with a *k* value of 2 with results compared to observed specimen sex, and optimum cluster number was assessed using the average silhouette (Rousseeuw, 1987) and gap statistic (Tibshirani et al., 2001) methods. Finally, Hartigans’ dip test was used to assess non-unimodality in the allometry-corrected shape dataset for the first 8 residual shape components (Maeckler, 2016).

Finally, we quantified shape variance in our dataset with the ‘morphol.disparity’ function in *geomorph* (Adams and Otárola-Castillo, 2013), for whole-skull and individual module shape data derived from both modularity analyses. Values were divided by the number of landmarks in each module to give a mean variance value that could be compared across different modules, and was repeated for allometry-corrected shape data.

## Results

Principal components analysis shows that the majority of shape variation (PC1, 57.4%) involves a change from narrow-span, small-horned shape at maximum value of PC1 to broad-span, large-horned shape at its minimum, with some corresponding relative increase in skull width around the orbits. The distribution of specimens on PC1 also suggests that sex is a discriminating factor, with female specimens (red) towards the positive end and male specimens (blue) towards the negative. PC2 (11.9% of total shape variation) shows mainly a change in horn shape, from curved horns that are flattened at the base, to more rounded horns which have a deep, pronounced boss at the contact with the skull.

The MANOVA performed on the raw shape data found a significant difference in shape between male and females in all specimens (n = 70, R^2^ = 0.21, *p* = 0.001), and when specimens of uncertain sex were removed (n = 54, R^2^ = 0.24, *p* = 0.001). Similarly, male and female centroid size was found to be significantly different (n = 70, F = 42.36, *p* = <0.001; Supplementary Fig. S4), including when analysing horns alone (F = 62.27, *p =* 0.001) and whole skull minus horns (F = 35.38, *p* = 0.001). The k-means cluster analysis performed reasonably well in identifying specimen groups, correctly assigning 81% of male and 89% of female specimens in raw shape data, and 72% of males and 93% of females in centroid size data (Supplementary Fig. S5). When optimum cluster number was assessed, however, results were mixed. The average silhouette method returned an optimum cluster number of 2 for the entire dataset, but also for male-only and female-only subsets of the sample (Supplementary Fig. S6). The Gap statistic method found no support for clustering in either the entire dataset, or male-only or female-only datasets. The dip test for non-unimodality was unable to identify non-unimodality in either shape data in the first eight PCs (Supplementary Table S5) or centroid size (D = 0.042, *p* = 0.48).

Allometry was found to have a strong and significant effect across the whole skull (n = 70, R^2^ = 0.38, *p* = 0.001). The difference in allometric slope between males and females was found to be non-significant (R^2^ = 0.01, *p* = 0.635), suggesting shared allometric trajectories. Moreover, no significant difference in shape was found between males and females when corrected for allometry (n = 70, R^2^ = 0.02, *p* = 0.206), suggesting that differences in shape between sexes is an artefact of size. This finding is further supported by the results of the k-means cluster analysis on allometry-corrected shape data, which correctly assigned 48% of female and 56% of male specimens, values no better than random group assignment (Supplementary Fig. S5). The results of the optimum cluster analyses echoed the uncorrected shape data, with the average silhouette method finding a 2-cluster optimum for the entire dataset, and the gap statistic method finding no support for clusters in the allometry-corrected dataset (Supplementary Fig. S6).

The maximum likelihood analysis of modularity hypotheses returned strongest support for the 12-modules hypothesis, of each bone plus the horn as an individual module (Fig. 2). ML approaches are known to preferentially support highly-paramaterised datasets, so we followed previous studies (Bardua et al. 2019a, Knapp et al., 2021) and examined the between-element correlation values to determine if any elements could reasonably be merged into a single module. Using a correlation value threshold of within 0.1 of either connected element we merged three pairs of elements (frontal and parietal, occipital and sphenoid, and premaxilla and maxilla), resulting in a 9-module configuration. The compare.CR analysis returned strongest support for a more integrated 3-module hypothesis (face, cranium and horns). Both modularity analyses thus supported weak integration of the horns with the rest of the skull.

**Figure 1:**
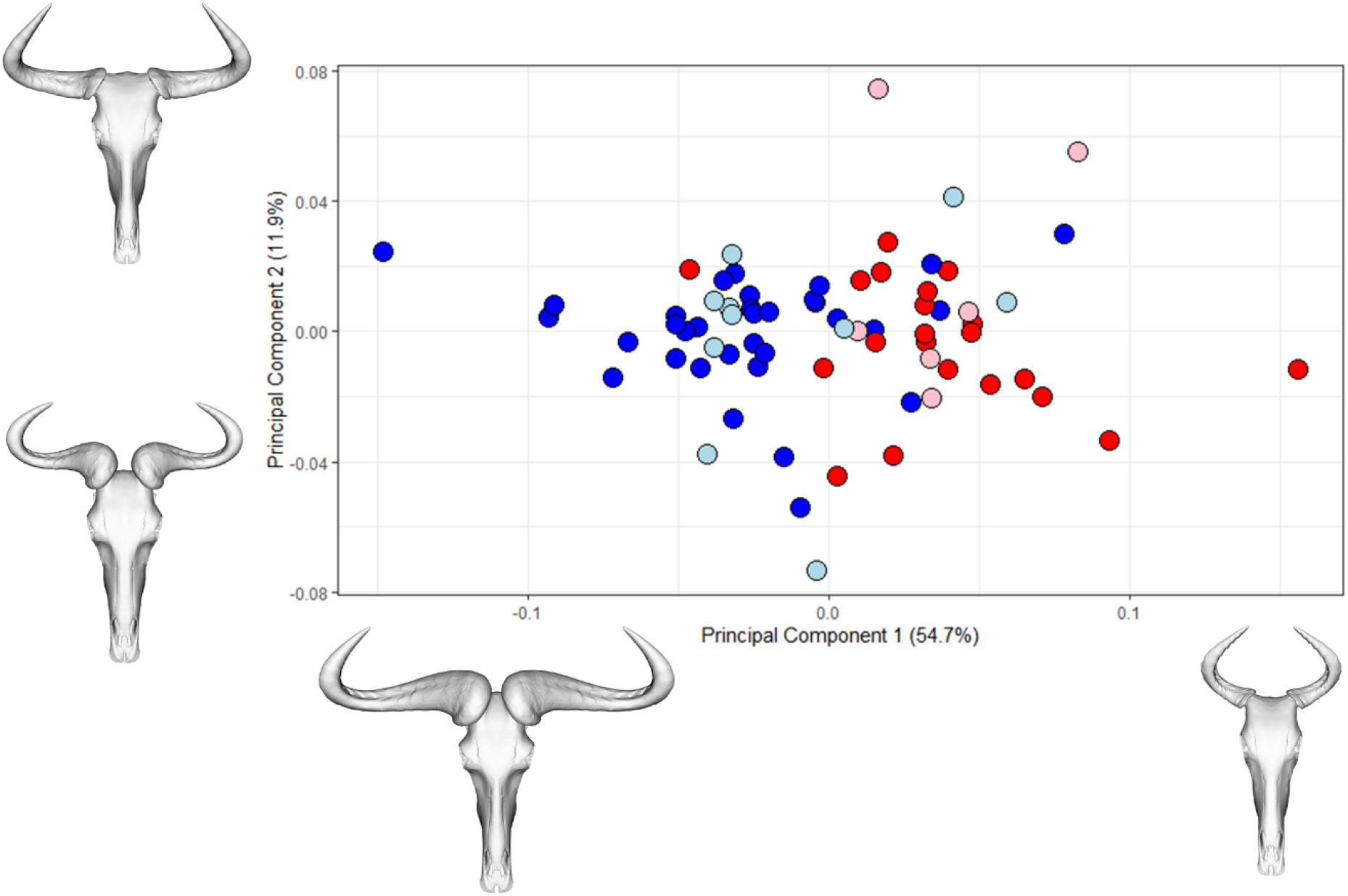
PCA of whole skull shape in *C. taurinus*, for first two principal components. Points are coloured according to sex (**dark blue**: male; **light blue**: probable male; **red**: female; **pink**: probable female). Projected skull shapes are shown in dorsal view along PC axes, representing extreme maximum and minimum shapes for each PC.

**Figure 2:**
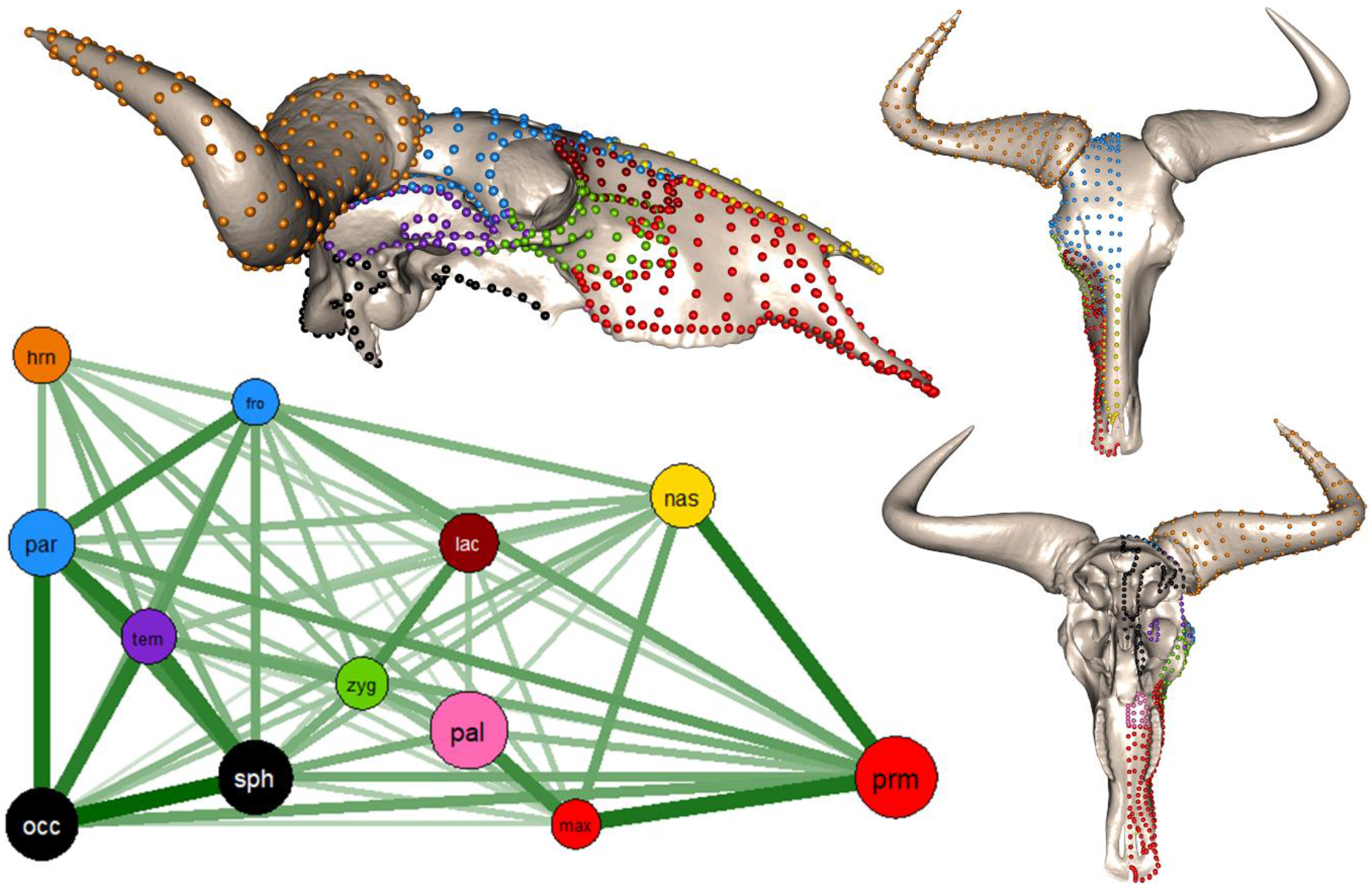
Network plot of EMMLi-derived covariance between skull elements, and corresponding landmarks. Circles on network plot represent separate bones and green lines represent covariance values between different bones. Size of circles and weight of lines represent intra- and inter element covariance values respectively. Landmarks on meshes are coloured according to network plot; matching colours indicate merged modules. Bone abbreviations are **hrn**: horn; **fro**: frontal; **par**: parietal; **lac**: lacrimal; **tem**: temporal; **zyg**: zygomatic; **occ**: occipital; **sph**: sphenoid; **pal**: palatine; **max**: maxilla; **prm**: premaxilla.

The shape of all phenotypic modules were found to be significantly correlated with size, with effect size ranging from 0.1 (lacrimal) to 0.44 (horn). The allometric slope of the horn was significantly higher than that of any other module (*p* = 0.001), in analyses of both shape and centroid size (Fig. 3), but the difference in allometric slope between male and female horn shape was found to be non-significant (R^2^ = 0.01, *p* = 0.635). The k-means clustering analysis performed on individual modules produced mixed results in identifying sex for most modules, correctly assigning more than 75% of specimens in for premaxilla/maxilla (83%), horn (86%) and face (81%; Supplementary Table S8). As with whole-skull shape data, k-means clustering performed poorly on allometry-corrected modules, not accurately assign any sex to a greater degree of accuracy than 72% (male face, Supplementary table S7). Of all phenotypic modules, the horns had the highest mean shape variance, both for raw and allometry-corrected data (Fig. 4).

**Fig. 3:**
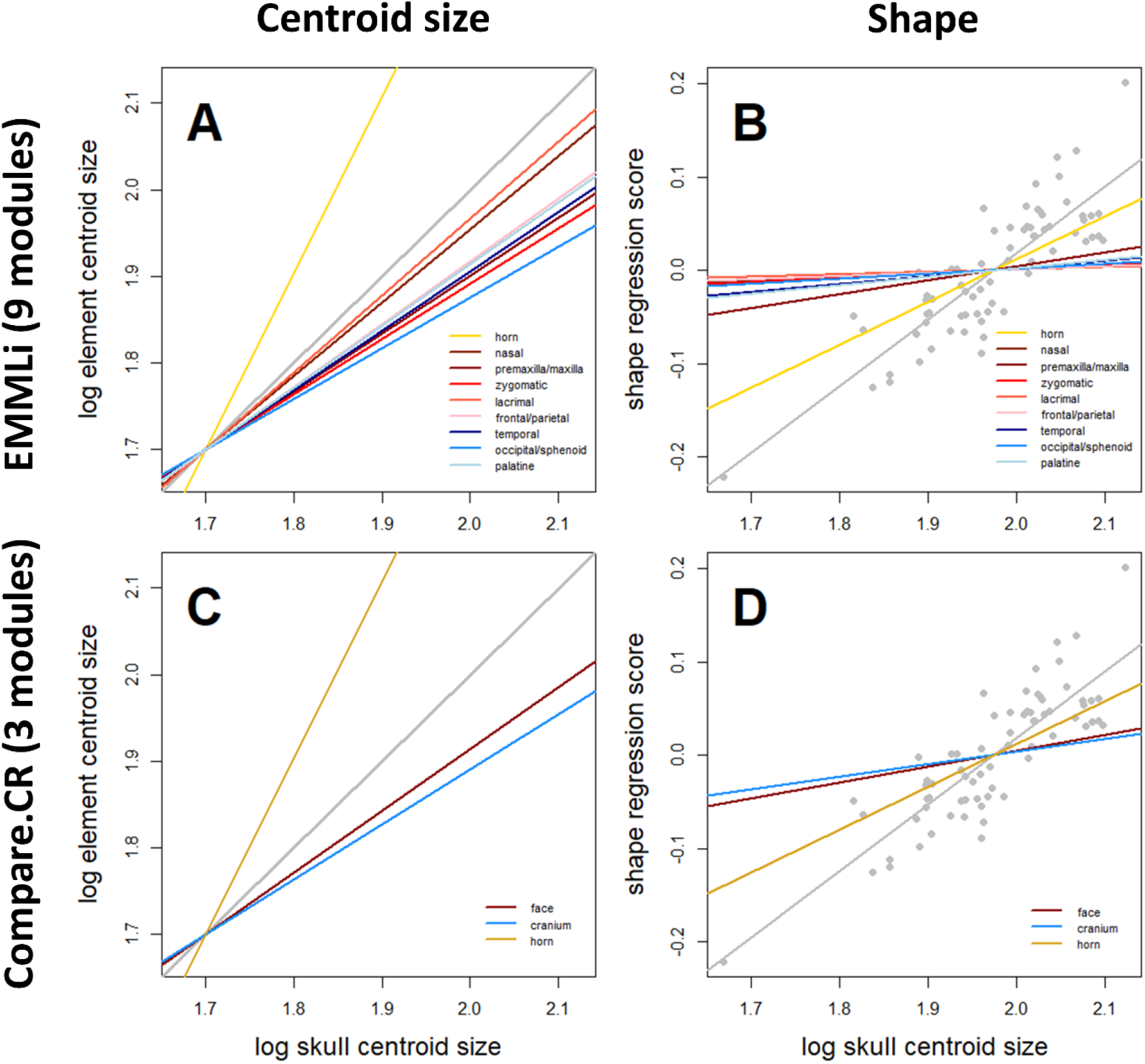
Allometry analyses of separate modules by centroid size (A and C) and shape (B and D). Slopes are coloured according to individual modules. Grey lines in each plot correspond to whole-skull. Plots correspond to 9-module model (A and B) derived from EMMLi modularity analysis, and 3-module model (C and D) derived from compare.CR analysis.

**Figure 4:**
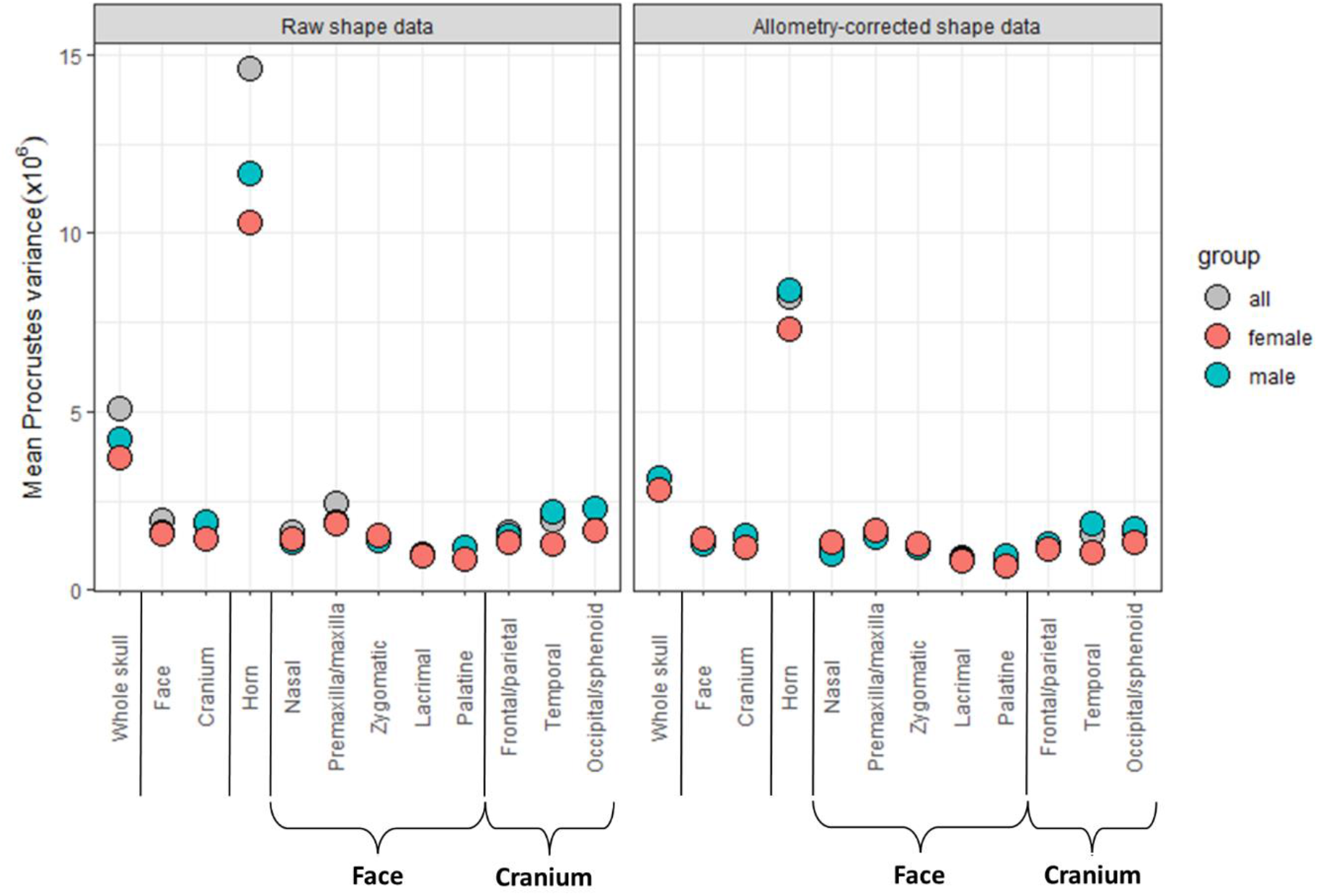
Per-module morphological variance. Shown are the mean Procrustes variance for whole skull and individual module for raw (left plot) and allometry-corrected (right plot) shape data. All modules from both *EMMLi* and *compare*.*CR* analyses are included, with subdivisions of the ‘face’ and ‘cranium’ module identified in the *compare*.*CR* analysis indicated. Points are coloured by sex.

## Discussion

Sexual selection is predicted to have consistent and detectable effects on the morphology of secondary sexual traits, and confirming these predictions in extant taxa, for which we have detailed information, is an important step in detecting it in extinct taxa. In this study we have shown that the sexually selected horns of the blue wildebeest, *C. taurinus*, fit hypothesised patterns of growth and variation, supporting the claim that secondary sexual traits may be readily detectable using morphology alone. Of the four hypotheses outlined in the introduction, three are supported by this study. First, skull shape differs significantly between sexes in *C. taurinus*; however, this difference is not significant after correcting for allometry. Second, the skull of *C. taurinus* has a modular structure, with the horns forming an internally integrated module which is weakly integrated with the remainder of the skull. Finally, the horns of *C. taurinus* show significant correlation with size and change shape at a higher rate than any other skull element. However, the fourth hypothesis, that the allometric trajectories of skull shape differ significantly between sexes is not supported. Our findings thus suggest that sexual dimorphism in *C. taurinus* is due to differences in size between sexes, with shape differences solely reflecting size differences due to allometry. Crucially, separating sexes within this dataset is impossible to verify without prior knowledge of sex, even with an exaggerated sexually selected trait which has been demonstrated to be sexually dimorphic. This issue is likely to apply widely unless sexual dimorphism is either in the form of presence/absence of sexually selected traits, or in extreme size dimorphism.

Intrasexual competition for mates often results in selection for larger size in the competing sex because larger individuals are physically dominant, and this may lead to the evolution of strong sexual dimorphism in size depending on the magnitude of this difference in selection (Tidière et al., 2017). Weapons to aid in physical competition, such as horns, may also evolve in the competing sex, and the presence of these traits within a population is sometimes used to infer sexual selection (Knell and Sampson, 2011). The evolution of strong positive allometry of these traits is consequently expected to evolve as a signal to amplify size and thus dominance (O’Brien et al., 2018). Our study supports this prediction with significant correlation of shape with size across the skull of *C. taurinus*, and the horns showing significantly higher rates of size and shape change than other skull elements. The increased variance of the horns compared to other skull elements, even when corrected for allometry, is another prediction of secondary sexual traits supported by our study. This is thought to be a result of relaxed functional constraints on the form of secondary sexual traits compared with other traits (O’Brien et al., 2018).

In some circumstances, secondary sexual traits may be expressed in the non-competing sex, but for other reasons (Tobias et al., 2012), and it might be expected that the degree of dimorphism in form may be amplified in such traits because of different selective pressures on each sex. When assessing sexual dimorphism in the dataset, however, our results were mixed. Although shape was found to be significantly sexually dimorphic, this was not the case when the dataset was corrected for allometry. This shift can be explained by the identical allometric trajectories of the sexes in *C. taurinus*. Similarly, the k-means clustering analysis performed well in identifying sexes in raw shape data (84% accuracy), but when shape data was corrected for allometry it performed no better than random (53% accuracy). Furthermore, the two methods employed for assessing optimum cluster number gave contradictory results that were not affected by either correcting for specimen size, or by removing either sex from the dataset entirely. These results have important implications for detecting sexual dimorphism; dimorphism can be strongly dependent on size, and allometry can act to magnify shape dimorphism between sexes.

Further complicating studies of sexual dimorphism, methods for detecting dimorphism, even in a large sample, have limitations. For example, similar to previous analyses of dimorphism, Hartigans’ dip test was particularly unsuccessful at detecting non-unimodality in this study (Mallon, 2017). Our dataset fits the recommended criteria outlined by Hone and Mallon (2017) for detectable dimorphism, in being a large sample size (>35 specimens) and a taxon with rapid growth to asymptotic size. Nevertheless, despite being seemingly segregated in a principal components analysis (Fig. 1), the overlap in shape and size between males and females was sufficient to mask robust recovery of dimorphism using the dip test method (Supplementary Table S5 and Fig. S4). Although *k*-means clustering performed well in distinguishing sexes in the raw data, the number of clusters is pre-determined by the user. When assessing sexual dimorphism, two clusters will therefore always be recovered, even when assessing a single-sex dataset. Furthermore, clustering accuracy can only be assessed with independent knowledge of specimen sex, which is not available for many fossil taxa (Mallon, 2017). In a dataset of adult specimens known to be significantly dimorphic, as in this study, this method works well in distinguishing sexes because of the difference in mean skull size between the two. In an ontogenetic study which captures the full range of morphologies from infant to fully-grown adult, it is more likely to separate juvenile and adult specimens by shape. *K*-means clustering analysis is therefore only appropriate where specimens are of a similar life stage, or when ontogenetic shape data are corrected for allometry.

Despite being an archetypal sexually selected structure in males, there is presently little agreement on the function of horns in female bovids (Tobias et al., 2012), even in well studied taxa such as *C. taurinus* (Estes, 2014). Predator defence, male mimicry, and genetic linkage to males (Estes, 2014) are the most frequently cited explanations for the presence of horns in female bovids. However, predator defence is seldom observed in *C. taurinus* and is often ineffective (Estes, 2014), although horns may act as a visual deterrent to predators in the open habitat in which this species tends to live (Stankowich and Caro, 2009). Male mimicry in this species is predicted to allow younger males to benefit from remaining in the maternal herd for longer (Estes, 2014), but there are two main problems with this hypothesis. Firstly, older males apparently have little problem in distinguishing and routinely evicting older males from herds of females and young (Estes, 2014). Secondly, it is unclear exactly how this would lead to the evolution of male mimicry by females rather than the evolution of female mimicry by males, given that males would benefit from remaining in a (majority) female herd. Horns in females may occur through genetic linkage, and similar expression of linked traits in both sexes is expected in this case. This is possible, but is not universal among bovids because the females of many bovid taxa are hornless (Packer, 1983). The probable cost of growing and maintaining such large and apparently costly traits suggests that their presence likely serves some adaptive function (Somjee, 2021), and it is probable that female horns are maintained by a combination of factors in *C. taurinus*.

Our results reveal that the skull of *C. taurinus* has a modular structure, with skull elements forming discrete phenotypic modules which are able to grow and vary with some independence from other elements. Although the two modularity analyses we used gave different results, both supported low integration of the horn with the rest of the skull. This is likely in secondary sexual traits because this allows horns to respond to selection with some degree of independence (Klingenberg, 2008). This can explain the strong positive allometric growth of this trait, and the considerably higher morphological variance of the horns, even when corrected for allometry (Fig. 4). Modularity may ultimately explain the evolution of a wide range of horn shapes across the bovid clade (Caro et al., 2003), and analysis of evolutionary modularity across Bovidae will help to support this prediction. Comparing modules across the entire skull is an important step in establishing the extreme growth of sexually selected traits, because it allows us to put the sexual trait in context with other aspects of anatomy and removes the tendency to focus on a single trait and introduce potential bias into the analysis.

Natural history collections have been shown to be biased towards male specimens, particularly in taxa with extreme secondary sexual traits (Cooper et al., 2019), and in this dataset males outnumber females by 43 to 27. Although strongly male-skewed, subdividing this dataset into equal numbers of each sex, or into individual sexes alone does not affect the overall results (Supplementary Tables S2 – S4). Historical collecting biases towards larger ‘trophy’ specimens may have the effect of creating distinct peaks of the largest individuals of both sexes, and fewer smaller individuals, which may create an even more marked sexual dimorphism than found in a natural population by decreasing the overlap between sexes. Furthermore, the keratin horn sheath is known to vary in size across different taxa relative to the bony horn core it encloses (Bubinek and Bubinek, 1990), and measurements taken on the horn sheath may therefore further amplify horn allometry. It is therefore likely that in fossil or osteology specimens, where soft tissues such as keratin are not preserved, that the effects of allometry and dimorphism will be less pronounced than in the specimens used here.

Sexual selection appears to have driven the evolution of both horns and sexual size dimorphism in *C. taurinus*. Our study has shown that the horns of *C. taurinus* displays patterns of growth and variation typically found in secondary sexual traits (O’Brien et al., 2018) in both sexes, despite strong sexual selection operating only in males. Although sexes are significantly different in size and shape, sexual dimorphism in the skull shape of *C. taurinus* appears to be a product of allometry. Our findings show that contrary to some previous claims (Knell and Sampson, 2011; Borkovic and Russell, 2014), determining dimorphism is not vital in detecting the trace of sexual selection in the skull of *C. taurinus*. Both sexes follow identical patterns of growth and variation across the skull, and are separated only by size. These results reflect attempts to recover dimorphism in extinct taxa where sex is not known *a priori* (Mallon, 2017; Knapp et al., 2021). Reproductive biology, life history and intensity of sexual selection may all affect the diversity, magnitude of sexual dimorphism and relative growth of secondary sexual traits, but the basic effects of sexual selection on morphology are likely to remain detectable to some extent. Ultimately, our findings suggest that identifying the patterns of growth and variation expected of sexually selected traits may be sufficient to identify secondary sexual traits using morphology alone, without the need to identify sexes through dimorphism.

## Supporting information

Additional figures and analyses summaries

## Acknowledgements

We are thankful to Roberto Portela Miguez and Natalie Cooper (NHM) for access to specimens, and Faisal Bibi (MfN) for providing additional specimen scans. We also thank Anjali Goswami for providing valuable feedback and comments on the manuscript.

## Author contributions

AK and CG designed the study. AK created scans of specimens. CG landmarked all specimens. AK and CG analysed data and wrote the manuscript.

## References

Adams DC and Otárola-Castillo E (2013). Geomorph: and R package for the collection and analysis of geometric morphometric shape data. Methods in Ecology and Evolution. 4; 393 –399

AgiSoft PhotoScan Professional (Version 1.4.3) (Software). (2019). Retrieved from http://www.agisoft.com/downloads/installer/.

Andersson M (1994). Sexual Selection. Princeton University Press, Princeton, NJ.

Arbour JH and Brown CM (2014). Incomplete specimens in geometric morphometric analyses. Methods in Ecology and Evolution. 5: 16–26. doi: 10.1111/2041-210X.12128.

Bardua C, Wilkinson M, Gower DJ, Sherratt E and Goswami A (2019). Morphological evolution and modularity of the caecilian skull. BMC Evolutionary Biology. doi:10.1186/s12862-018-1342-7.

Bardua C, Felice RN, Watanabe A, Fabre AC and Goswami A (2019). A practical guide to sliding and surface semilandmarks in morphometric analyses. Integrative Organismal Biology. Obz016, doi: 10.1093/iob/obz016.

Bonduriansky R (2007). Sexual selection and allometry: A critical reappraisal of the evidence and ideas. Evolution. 61; 838 – 849

Borkovic B and Russell A (2014). Sexual selection according to Darwin: A response to Padian and Horner’s interpretation. C. R. Palevol. 13: 701 – 707

Bubenik GA and Bubinek AB (eds). Horns, Pronghorns and Antlers. Springer-Verlag, New York. 1990.

Caro TM, Graham CM, Stoner CJ and Flores MM (2003). Correlates of horn and antler shape in bovids and cervids. Behav. Ecol. Sociobiol. 55; 32 – 41

Castelló JR (2016). Bovids of the World. Princeton University Press, Princeton and Oxford.

Cooper N, Bond AL, Davis JL, Portela Miguez R, Tomsett L, Helgen KM (2019) Sex biases in bird and mammal natural history collections. Proceedings of the Royal Society B: Biological Sciences, 286 (1913) : 20192025 –20192025. doi: 10.1098/rspb.2019.2025.

Darwin C (1871). The descent of man and selection in relation to sex. London, UK: John Murray.

Estes RD (2014). The Gnu’s World. University of California Press, Berkeley and Los Angeles, California.

Evans KM, Bernt MJ, Kolmann MA, Ford KL and Albert JS (2018). Why the long face? Static allometry in the sexually dimorphic phenotypes of Neotropical electric fishes. Zool. Jour. Linn. Soc. doi: 10.1093/zoolinnean/zly076.

Goswami A and Finarelli JA (2016). EMMLi: A maximum likelihood approach to the analysis of modularity. Evolution. doi: 10.1111/evo.12956.

Goswami A, Watanabe A, Felice RN, Bardua C, Fabre AC and Polly PD (2019). High-density morphometric analysis of shape and integration: the good, the bad, and the not-really-a-problem. Integrative and Comparative Biology. https://doi.org/10.1093/icb/icz120

Hartigan JA and Wong M A (1979). Algorithm AS 136: A K-means clustering algorithm. Applied Statistics, 28, 100–108. doi: 10.2307/2346830.

Hone DWE and Mallon JC. (2017). Protracted growth impedes the detection of sexual dimorphism in non-avian dinosaurs. Palaeontology. 1 –11. doi: 10.1111/pala.12298

Janicke T, Ritchie MG, Morrow EH, Marie-Orleach L (2018). Sexual selection predicts species richness across the animal kingdom. Proc. R. Soc. B 285, 20180173. (xDoi:10.1098/rspb.2018.0173)

Klingenberg CP (2008). Morphological integration and developmental modularity. Annu. Rev. Ecol. Syst. 39; 115 – 132.

Knapp A, Knell RJ and Hone DWE (2021). Three-dimensional geometric morphometric analysis of the skull of Protoceratops andrewsi supports a socio-sexual signalling role for the ceratopsian frill. Proc. R. Soc. B. 288: 20202938. https://doi.org/10.1098/rspb.2020.2938.

Knell RJ and Sampson S (2011). Bizarre structures in dinosaurs: species recognition or sexual selection? A response to Padian and Horner. Journal of Zoology. 283; 18 – 22.

Knell RJ, Naish D, Tompkins JL, and Hone DWE (2012). Sexual selection in prehistoric animals: detection and implications. TREE. 28; 38 – 47.

Losos JB (2011). Convergence, adaptation and constraint. Evolution. 65 ; 1827 – 1840.

Maechler M (2016). diptest: Hartigans’ Dip Test Statistic for Unimodality - Corrected. R package version 0.75-7. https://CRAN.R-project.org/package=diptest.

Mallon JC (2017). Recognizing sexual dimorphism in the fossil record: lessons from nonavian dinosaurs. Paleobiology 43, 495–507. (doi:10.1017/pab.2016.51)

Martínez-Ruiz C and Knell RJ (2016). Sexual selection can both increase and decrease extinction probability: reconciling demographic and evolutionary factors. J. Anim. Ecol. 86, 117–127. (doi:10.1111/1365-2656.12601)

Mitteroecker P, Gunz P, Bernhard M, Schaefer K and Bookstein FL (2004). Comparison of cranial ontogenetic trajectories among great apes and humans. Journal of Human Evolution. 46; 679 – 698 doi:10.1016/j.jhevol.2004.03.006

O’Brien DM, Allen CE, Van Kleeck MJ, Hone DWE, Knell RJ, Knapp A, Christiansen S and Emlen DJ (2018). On the evolution of extreme structures: static scaling and the function of sexually selected signals. Animal Behaviour. 144; 95 – 108.

Packer C (1983). Sexual dimorphism: the horns of African antelopes. Science. 221: 1191 – 1193.

R Core Team (2021). R: A language and environment for statistical computing. R Foundation for Statistical Computing, Vienna, Austria. URL https://www.R-project.org/.

Ritchie MG (2007). Sexual selection and speciation. Annu. Rev. Ecol. Evol. Syst. 38, 79–102.

Rodríguez RL and Eberhard WG (2019). Why the static allometry of sexually—selected traits is so variable: The importance of function. Integrative and Comparative Biology. Doi:10.1093/icb/icz039.

Rousseeuw PJ (1987). Silhouettes: a graphical aid to the interpretation and validation of cluster analysis.. Journal of Computational and Applied Mathematics. 20: 53–65.

Schlager S (2017). Morpho and Rvcg - Shape Analysis in R. In Zheng G, Li S, Szekely G (eds.), Statistical Shape and Deformation Analysis, 217–256. Academic Press. ISBN 9780128104934.

Somjee U (2021). Positive allometry of sexually selected traits: Do metabolic maintenance costs play an important role? BioEssays. https://doi.org/10.1002/bies.202000183.

Stankowich T and Caro T (2009). Evolution of weaponry in female bovids. Proc. R. Soc. B. 276; 4329 – 4334.

Stratovan Checkpoint v. 2020.10.13.0859. Stratovan Corporation. Available from https://www.stratovan.com/products/checkpoint.

Tibshirani R, Walther G and Hastie T (2001). Estimating the number of clusters in a dataset via the gap statistic. J. R. Statist. Soc. B. 63: 411–423.

Tidière M, Lemaître JF, Pélabon C, Giminez O and Gaillard JM (2017). Evolutionary allometry reveals a shift in selection pressure in male horn size. J. Evol. Bio. https://doi.org/10.1111/jeb.13142

Tidière M, Gaillard JM, Garel M, Lemaître JF, Toïgo C and Pélabon C (2020). Variation in the ontogenetic allometry of horn length in bovids along a body mass continuum. Ecology and Evolution. 10: 4104 – 4114.

Tobias JA, Montgomerie R and Lyon BE (2012). The evolution of female ornaments and weaponry: social selection, sexual selection and ecological competition. Phil. Trans. R. Soc. B. 367; 2274 – 2293.

West-Eberhard MJ (1979). Sexual selection, social competition and evolution. Amer. Phil. Soc. 123; 222 – 234.

Zelditch ML, Swiderski DL, Sheets HD and Fink WL (2004). Geometric Morphometrics for Biologists: A Primer. Elsevier Inc.

Zelditch ML and Goswami A (2021). What does modularity mean? Evolution and Development. 23: 377 – 403. DOI: 10.1111/ede.12390

