## Additional figures and analyses summaries for "Sexual selection leads to positive allometry but not sexual dimorphism in the expression of horn shape in the blue wildebeest, *Connochaetes taurinus*"

### Supplementary data

#### Note on sample and additional analyses

The sample consists of 75 *C. taurinus* skulls; 47 males and 28 females. Of these specimens, 5 were too severely damaged to place surface semilandmarks and so were excluded from the surface semilandmark analysis (i.e. **patched specimens**), and were analysed only with anatomical landmarks and semilandmark curves (i.e. **curves only**). Sex was known for certain (i.e., information was included on specimen labels) for 36 males and 22 female specimens. The sex of the remaining specimens was estimated by the authors and marked as 'probable male' or 'probable female.' The dataset was subdivided to include only specimens of certain sex (i.e. '**strict**' as opposed to '**all**,' which also included specimens with estimated sex), and analyses were rerun on the '**strict**' subset. In addition, all analyses were run on individual sexes (**Males only** and **Females only**), for all specimens and only those of known sex. Because males outnumber females, the largest males were removed from the dataset to equalise the number of each sex ('**equal sample**'), for both the '**all**' and '**strict**' subsets.

Specimens in the dataset represent all 5 subspecies of *C. taurinus*: *C. t. taurinus* (n = 36), *C. t. johnstoni* (n = 4), *C. t. mearnsi* (n = 4), *C. t. albojubatus* (n = 6) and *C. t. cooksoni* (n = 3). In addition, 15 specimens are of unknown subspecies. To account for possible differences in shape between these subspecies a MANOVA was applied to the entire dataset with subspecies as the independent grouping variable, which did not reveal a significant difference between any group (Supplementary Data Table S3). Nevertheless, all analyses were repeated solely on specimens of the best-represented subspecies, *C. t. taurinus* (n = 36). These analyses are reported as '**subspecies analysis**' in the results tables below. The results of these analyses did not differ from our analysis of the entire dataset. Subspecies of *C. taurinus* are mainly distinguished by colouration and median body weight (Castelló, 2016), with horn shape being somewhat distinct in the Western White wildebeest, *C. t. mearnsi* (Estes, 2014; Castelló, 2016). Differences in horn shape that are independent of size are seen along PC2 of Fig. 1 of the main text, but are not sufficient to distinguish subspecies in this dataset. It may be the case that increasing the sample sizes of different subspecies may reveal more reliable differences between groups in this way.

**Table S1: Specimen data.** F = female, M = male, F_p = probable female, M_p = probable male.

| specimen | species | subspecies | sex |
|---|---|---|---|
| NHM1842.4.11.10 | connochaetes_taurinus | C. t. taurinus | M |
| NHM1863.7.7.6 | connochaetes_taurinus | C. t. johnstoni | F |
| NHM1893.6.20.3 | connochaetes_taurinus | C. t. albojubatus | F |
| NHM1899.6.29.5 | connochaetes_taurinus | NA | M |
| NHM1900.3.18.14 | connochaetes_taurinus | NA | M |
| NHM1900.3.18.15 | connochaetes_taurinus | NA | F |
| NHM1902.11.18.3 | connochaetes_taurinus | C. t. albojubatus | M |
| NHM1902.11.18.4 | connochaetes_taurinus | NA | F |
| NHM1907.4.12.1 | connochaetes_taurinus | C. t. taurinus | F_p |
| NHM1907.10.4.8 | connochaetes_taurinus | NA | M |
| NHM1908.3.17.4 | connochaetes_taurinus | C. t. taurinus | M |
| NHM1910.9.26.1 | connochaetes_taurinus | NA | M_p |
| NHM1910.9.26.2 | connochaetes_taurinus | NA | M_p |
| NHM1910.9.26.3 | connochaetes_taurinus | NA | F_p |
| NHM1917.6.26.4 | connochaetes_taurinus | NA | F |
| NHM1919.7.15.118 | connochaetes_taurinus | NA | M |
| NHM1919.7.15.119 | connochaetes_taurinus | NA | M |
| NHM1919.7.15.122 | connochaetes_taurinus | NA | M_p |
| NHM1920.3.21.2 | connochaetes_taurinus | NA | M_p |
| NHM1921.7.18.18 | connochaetes_taurinus | C. t. taurinus | F |
| NHM1925.12.4.232 | connochaetes_taurinus | C. t. taurinus | M |
| NHM1927.8.16.5 | connochaetes_taurinus | C. t. taurinus | M |
| NHM1927.8.16.6 | connochaetes_taurinus | C. t. taurinus | M |
| NHM1930.12.3.5 | connochaetes_taurinus | C. t. taurinus | M |
| NHM1930.12.3.6 | connochaetes_taurinus | C. t. taurinus | M |
| NHM1931.2.1.15 | connochaetes_taurinus | C. t. taurinus | M |
| NHM1931.2.1.16 | connochaetes_taurinus | C. t. taurinus | F |
| NHM1931.2.1.17 | connochaetes_taurinus | C. t. taurinus | M |
| NHM1931.2.1.18 | connochaetes_taurinus | C. t. taurinus | M |
| NHM1932.6.6.27 | connochaetes_taurinus | NA | M |
| NHM1932.6.6.28 | connochaetes_taurinus | NA | F |
| NHM1932.6.6.29 | connochaetes_taurinus | NA | F |
| NHM1932.9.1.205 | connochaetes_taurinus | C. t. taurinus | M |
| NHM1934.11.1.13 | connochaetes_taurinus | C. t. taurinus | M |
| NHM1934.11.1.14 | connochaetes_taurinus | C. t. taurinus | F |
| NHM1935.3.16.3 | connochaetes_taurinus | C. t. taurinus | M |
| NHM1935.9.1.323 | connochaetes_taurinus | C. t. taurinus | M |
| NHM1935.9.1.324 | connochaetes_taurinus | C. t. taurinus | F |
| NHM1935.12.14.3 | connochaetes_taurinus | C. t. albojubatus | M_p |
| NHM1936.3.30.15 | connochaetes_taurinus | C. t. taurinus | M |
| NHM1936.3.30.16 | connochaetes_taurinus | C. t. taurinus | F |
| NHM1938.7.8.16 | connochaetes_taurinus | C. t. taurinus | M |
| NHM1938.7.8.17 | connochaetes_taurinus | C. t. taurinus | F |
| NHM1940.83 | connochaetes_taurinus | C. t. johnstoni | M_p |
| NHM1962.807 | connochaetes_taurinus | C. t. taurinus | M |
| NHM1962.808 | connochaetes_taurinus | C. t. taurinus | M |
| NHM1962.809 | connochaetes_taurinus | C. t. taurinus | M |
| NHM1966.464 | connochaetes_taurinus | C. t. taurinus | M |
| NHM1966.465 | connochaetes_taurinus | C. t. taurinus | F |
| NHM1966.466 | connochaetes_taurinus | NA | M |
| NHM1966.467 | connochaetes_taurinus | C. t. taurinus | F_p |
| NHM1966.468 | connochaetes_taurinus | NA | F |
| NHM1966.469 | connochaetes_taurinus | C. t. taurinus | F |
| NHM1966.47 | connochaetes_taurinus | C. t. taurinus | M |
| NHM1966.471 | connochaetes_taurinus | C. t. taurinus | F |
| NHM1966.472 | connochaetes_taurinus | C. t. taurinus | F |
| NHM1966.839 | connochaetes_taurinus | C. t. cooksoni | M |
| NHM1966.84 | connochaetes_taurinus | C. t. cooksoni | M |
| NHM1966.841 | connochaetes_taurinus | C. t. taurinus | M |
| NHM1967.1327 | connochaetes_taurinus | C. t. taurinus | M |
| NHM1968.692 | connochaetes_taurinus | C. t. taurinus | F |
| NHM1968.693 | connochaetes_taurinus | C. t. taurinus | M |
| NHM1971.2128 | connochaetes_taurinus | NA | M |
| NHM1971.2129 | connochaetes_taurinus | NA | M_p |
| NHM1975.1150 | connochaetes_taurinus | C. t. mearnsi | F |
| NHM1975.1152 | connochaetes_taurinus | NA | F |
| NHM1975.1154 | connochaetes_taurinus | NA | F |
| NHM1986.1406 | connochaetes_taurinus | NA | M |
| NHM1986.1407 | connochaetes_taurinus | NA | F_p |
| NHM2004.510 | connochaetes_taurinus | NA | M_p |
| NHM2004.511 | connochaetes_taurinus | NA | M_p |
| NHMXXX1 | connochaetes_taurinus | NA | F_p |
| NHMXXX2 | connochaetes_taurinus | NA | M_p |
| ZMB31826 | connochaetes_taurinus | NA | F_p |
| ZMB31843 | connochaetes_taurinus | NA | M_p |

**Modularity tests**

**Skull elements**

1: Nasal

2: Premaxilla

3: Maxilla

4: Zygomatic

5: Lacrimal

6: Frontal

7: Parietal

8: Horn

9: Occipital

10: Temporal

11: Sphenoid

12: Palatine

**Modularity hypotheses tested with ML method (EMMLi).** Numbers in parenthesis correspond to skull elements listed above.

1. All bones separate modules (1:12); **12 modules**
2. **1.** Horn (8), **2.** all other bones (1:7, 9:12); **2 modules**
3. **1.** Face (1:5,12), **2.** cranium (6,7,9:11), **3.** horn (8); **3 modules**
4. **1.** Face (1:5,12), **2.** horn + cranium (6:11); **2 modules**
5. **1.** Nasal (1), **2.** Premaxilla + maxilla (2,3), **3.** zygomatic + lacrimal (4,5), **4.** frontal (6), **5.** parietal (7), **6.** horn (8), **7.** occipital + sphenoid (9,11), **8.** temporal (10), **9.** palatine (12); **9 modules**
6. **1.** Nasal (1), **2.** premaxilla + maxilla (2,3), **3.** zygomtic + lacrimal (4,5), **4.** frontal (6), **5.** parietal + occipital + sphenoid (7,9,11), **6.** horn (8), **7.** temporal (10), **8.** palatine (12); **8 modules**
7. **1.** Nasal (1), **2.** premaxilla + maxilla (2,3), **3.** zygomatic (4), **4.** lacrimal (5), **5.** frontal + parietal (6,7), **6.** horn (8), **7.** occipital + sphenoid (9,11), **8.** temporal (10), **9.** palatine (12); **9 modules**

**Table S2: Results of allometry analyses on shape data.**

| | Curves only | | | Patched specimens | | |
|---|---|---|---|---|---|---|
| | **n** | **$R^2$** | ***p*** | **n** | **$R^2$** | ***p*** |
| **M+F - All** | 75 | 0.22 | 0.001 | 70 | 0.38 | 0.001 |
| **M+F – All, equal sample** | 56 | 0.18 | 0.001 | 54 | 0.33 | 0.001 |
| **M+F - Strict** | 58 | 0.22 | 0.001 | 54 | 0.37 | 0.001 |
| **M+F – Strict, equal sample** | 44 | 0.20 | 0.001 | 42 | 0.36 | 0.001 |
| **Males only – All** | 47 | 0.14 | 0.001 | 43 | 0.25 | 0.001 |
| **Males only - Strict** | 36 | 0.14 | 0.001 | 33 | 0.25 | 0.001 |
| **Females only – All** | 28 | 0.15 | 0.001 | 27 | 0.24 | 0.001 |
| **Females only - Strict** | 22 | 0.15 | 0.007 | 21 | 0.31 | 0.002 |
| **Subspecies analysis (*C. t. taurinus* only)** | n/a | n/a | n/a | 36 | 0.37 | 0.001 |

**Table S3: Comparison of Allometric slopes by sex.**

| | Curves only | | | Patched specimens | | |
|---|---|---|---|---|---|---|
| | **n** | **$R^2$** | ***p*** | **n** | **$R^2$** | ***p*** |
| **All** | 75 | 0.01 | 0.366 | 70 | 0.01 | 0.635 |
| **All, equal samples** | 56 | 0.03 | 0.017 | 54 | 0.02 | 0.145 |
| **All, horns only** | | | | 70 | 0.003 | 0.848 |
| **Strict** | 58 | 0.01 | 0.464 | 54 | 0.01 | 0.605 |
| **Strict, equal samples** | 44 | 0.03 | 0.046 | 42 | 0.02 | 0.116 |
| **Subspecies analysis** | | | | 36 | 0.03 | 0.146 |

**Table S4: MANOVA analyses of differences in shape by sex and subspecies.** Significant results are shown in **bold.**

| | Curves only | | | Patched specimens | | |
|---|---|---|---|---|---|---|
| | **n** | **$R^2$** | ***p*** | **n** | **$R^2$** | ***p*** |
| **All** | 75 | 0.16 | **0.001** | 70 | 0.21 | **0.001** |
| **All, equal samples** | 56 | 0.12 | **0.001** | 54 | 0.15 | **0.001** |
| **All, allometry corrected** | 75 | 0.04 | **0.005** | 70 | 0.02 | 0.206 |
| **All, equal samples, allometry corrected** | 56 | 0.06 | **0.001** | 54 | 0.04 | 0.052 |
| **Strict** | 58 | 0.19 | **0.001** | 54 | 0.24 | **0.001** |
| **Strict, equal samples** | 44 | 0.17 | **0.001** | 42 | 0.19 | **0.001** |
| **Strict, allometry corrected** | 58 | 0.05 | **0.005** | 54 | 0.03 | 0.139 |
| **Strict, equal samples, allometry corrected** | 44 | 0.07 | **0.004** | 42 | 0.04 | 0.126 |
| **Subspecies analysis, all specimens** | | | | 53 | 0.11 | 0.107 |
| ***C. t. taurinus* subset, sex** | | | | 36 | 0.25 | **0.001** |

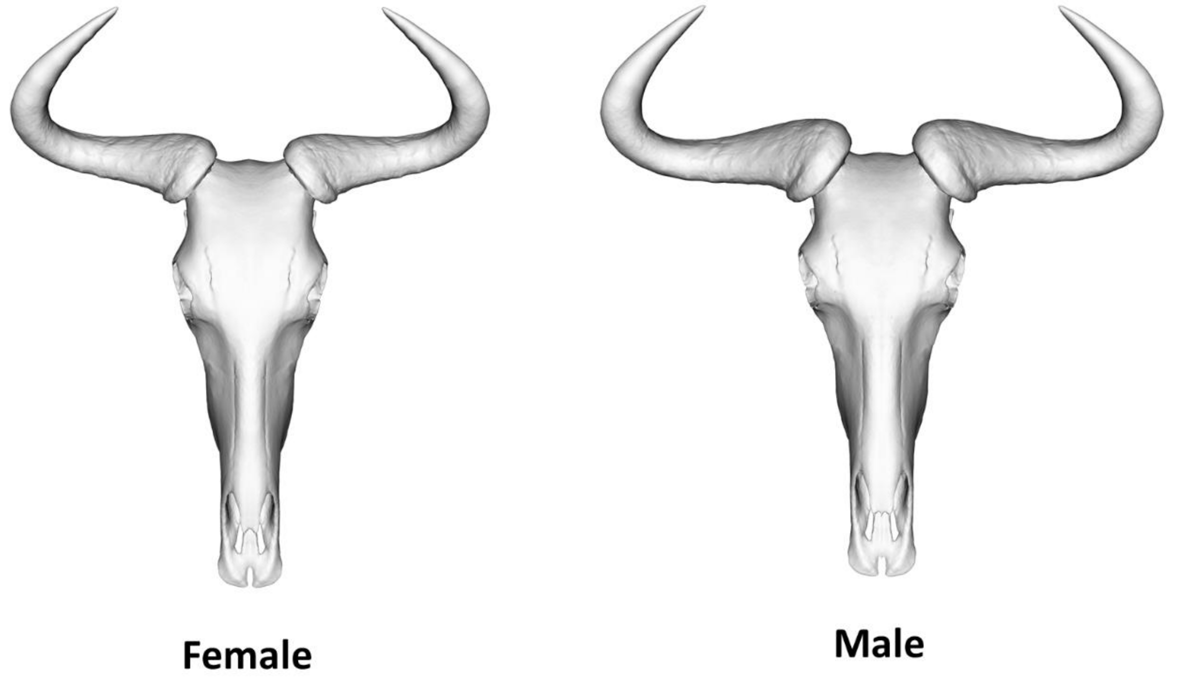


**Figure S1: Warped meshes representing mean shape of female (left) and male (right) *C. taurinus* skull.**

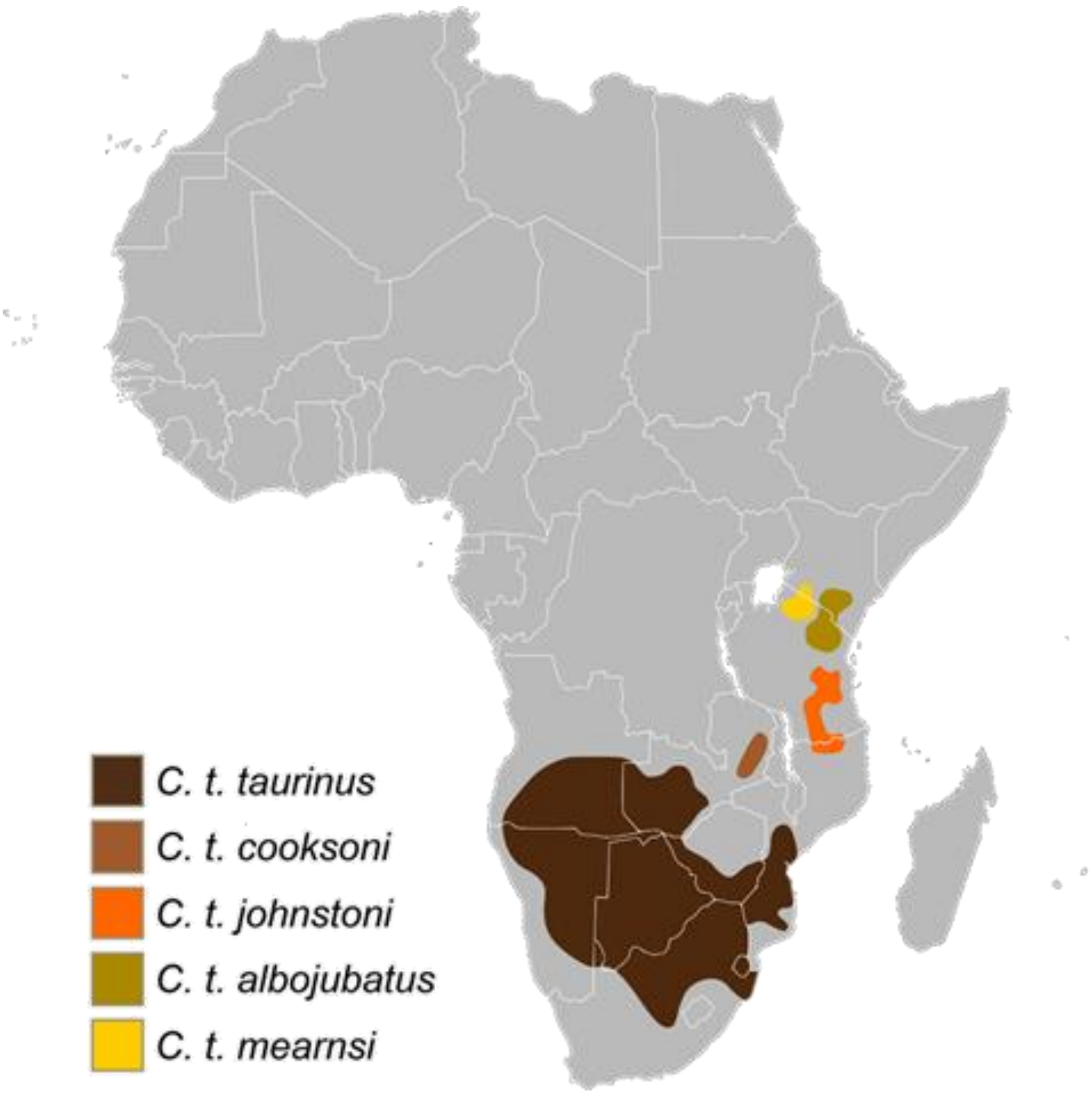


**Figure S2: Distribution of *C. taurinus* subspecies across Africa (licensed under CC BY-SA 3.0).**

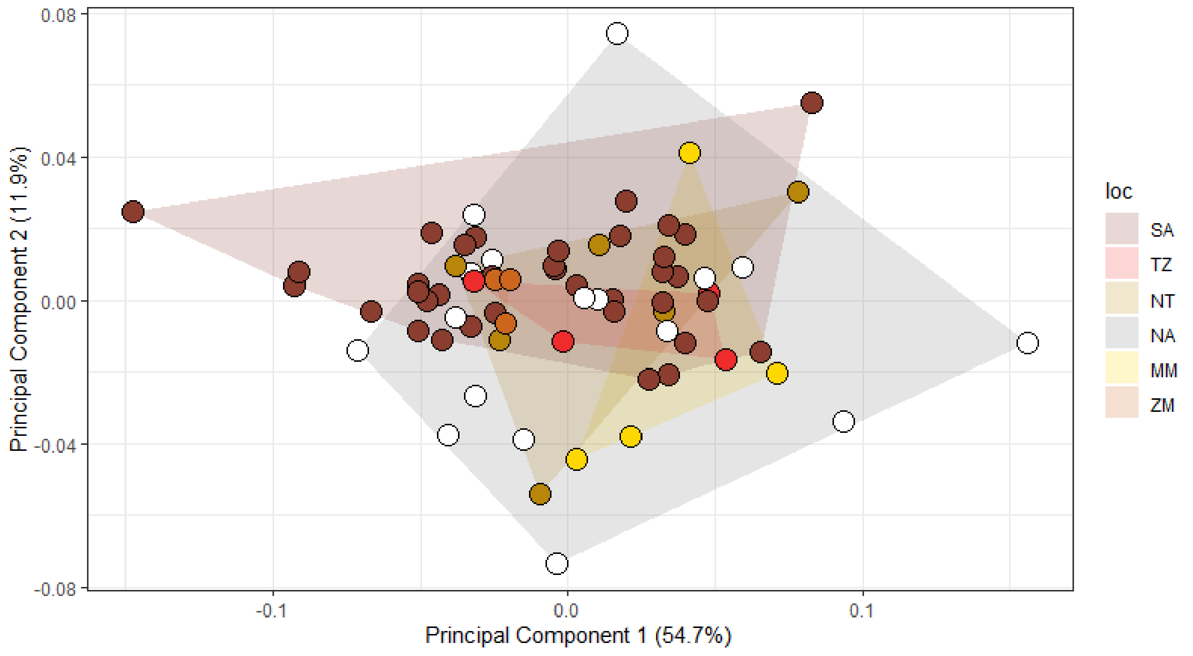


**Figure S3: PCA of specimens coloured by subspecies.** Convex hulls group subspecies, SA: *C. t. taurinus ;* TZ: *C. t. johnstoni*; NT: *C. t. albojubatus*; MM: *C. t. mearnsi*; ZM: *C. t. cooksoni;* NA: not known.

**Table S5: Results of dip tests performed on shape and size data.**

| | **Raw shape data** | | | **Allometry-corrected data** | | |
|---|---|---|---|---|---|---|
| | **PC proportion of variance (%)** | **D** | ***p*** | **PC proportion of variance (%)** | **D** | ***p*** |
| **Principal component 1** | 54.7 | 0.046 | 0.34 | 31.9 | 0.041 | 0.55 |
| **Principal component 2** | 11.9 | 0.029 | 0.98 | 18.4 | 0.044 | 0.38 |
| **Principal component 3** | 7.3 | 0.042 | 0.51 | 11.1 | 0.030 | 0.95 |
| **Principal component 4** | 4.3 | 0.039 | 0.65 | 6.0 | 0.024 | 1.00 |
| **Principal component 5** | 3.1 | 0.035 | 0.80 | 4.3 | 0.036 | 0.76 |
| **Principal component 6** | 2.6 | 0.029 | 0.98 | 4.1 | 0.034 | 0.86 |
| **Principal component 7** | 1.6 | 0.036 | 0.80 | 2.6 | 0.049 | 0.23 |
| **Principal component 8** | 1.5 | 0.034 | 0.83 | 2.1 | 0.032 | 0.89 |
| **Dip test, All, centroid size** | n/a | 0.042 | 0.48 | n/a | n/a | n/a |

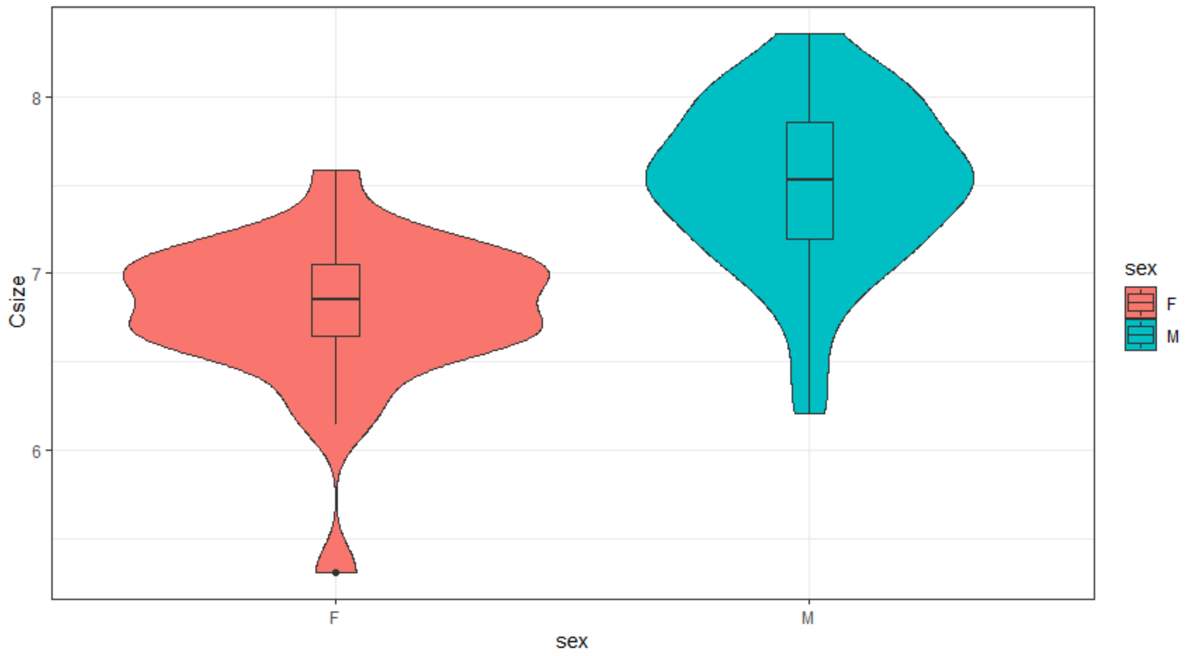


**Figure S4: Violin plots of centroid size distributions for females (F) and males (M).**

**Table S6: ANOVA of centroid size by sex**

| | Curves only | | Patched specimens | |
|---|---|---|---|---|
| | **F** | ***p*** | **F** | ***p*** |
| **All** | 34.2 | <0.001 | 42.36 | <0.001 |
| **Strict** | 30.72 | <0.001 | 40.55 | <0.001 |
| **All, horns only** | n/a | n/a | 62.27 | <0.001 |
| **All, skull minus horns** | n/a | n/a | 35.38 | <0.001 |

**Table S7: Results of shape allometry analysis of individual modules for all specimens**

| | EMMLi modules | | |
|---|---|---|---|
| | **n** | **Effect size** | ***p*** |
| **Nasal** | 70 | 0.21 | 0.004 |
| **Premaxilla/maxilla** | 70 | 0.31 | 0.004 |
| **Frontal/parietal** | 70 | 0.19 | 0.004 |
| **Zygomatic** | 70 | 0.15 | 0.004 |
| **Horn** | 70 | 0.44 | 0.004 |
| **Temporal** | 70 | 0.18 | 0.004 |
| **Lacrimal** | 70 | 0.10 | 0.004 |
| **Occipital/sphenoid** | 70 | 0.28 | 0.004 |
| **Palatine** | 70 | 0.28 | 0.004 |
| | **compare.CR modules** | | |
| **Face** | 70 | 0.27 | 0.004 |
| **Cranium** | 70 | 0.22 | 0.004 |
| **Horn** | 70 | 0.44 | 0.004 |

**Table S8: Results of k-means cluster analysis on whole-skull and individual module shape data, for raw and allometry-corrected shape data.** Results show percentage of individuals correctly assigned to group. Values above 75% are shown in bold.

| | Raw shape data | | | Allometry corrected shape data | | |
|---|---|---|---|---|---|---|
| | Males % correct | Females % correct | Total % correct | Males % correct | Females % correct | Total % correct |
| **Whole skull** | **81%** | **89%** | **84%** | 56% | 48% | 53% |
| **Nasal** | 72% | 74% | 73% | 58% | 63% | 60% |
| **Premaxilla/maxilla** | **84%** | **81%** | **83%** | 58% | 50% | 61% |
| **Frontal/parietal** | 65% | **89%** | 74% | 70% | 37% | 50% |
| **Zygomatic** | 60% | 52% | 57% | 52% | 53% | 53% |
| **Horn** | **79%** | **96%** | **86%** | 56% | 48% | 53% |
| **Temporal** | 72% | 48% | 63% | 51% | 52% | 51% |
| **Lacrimal** | 53% | 59% | 56% | 51% | 52% | 51% |
| **Occipital/sphenoid** | 65% | **85%** | 73% | 56% | 44% | 51% |
| **Palatine** | 65% | 70% | 67% | 51% | 60% | 54% |
| **Face (compare.CR)** | **81%** | **81%** | **81%** | 72% | 67% | 70% |
| **Cranium (compare.CR)** | 74% | 74% | 74% | 47% | 67% | 59% |
| **Centroid size, whole skull** | 72% | **93%** | **80%** | n/a | | |

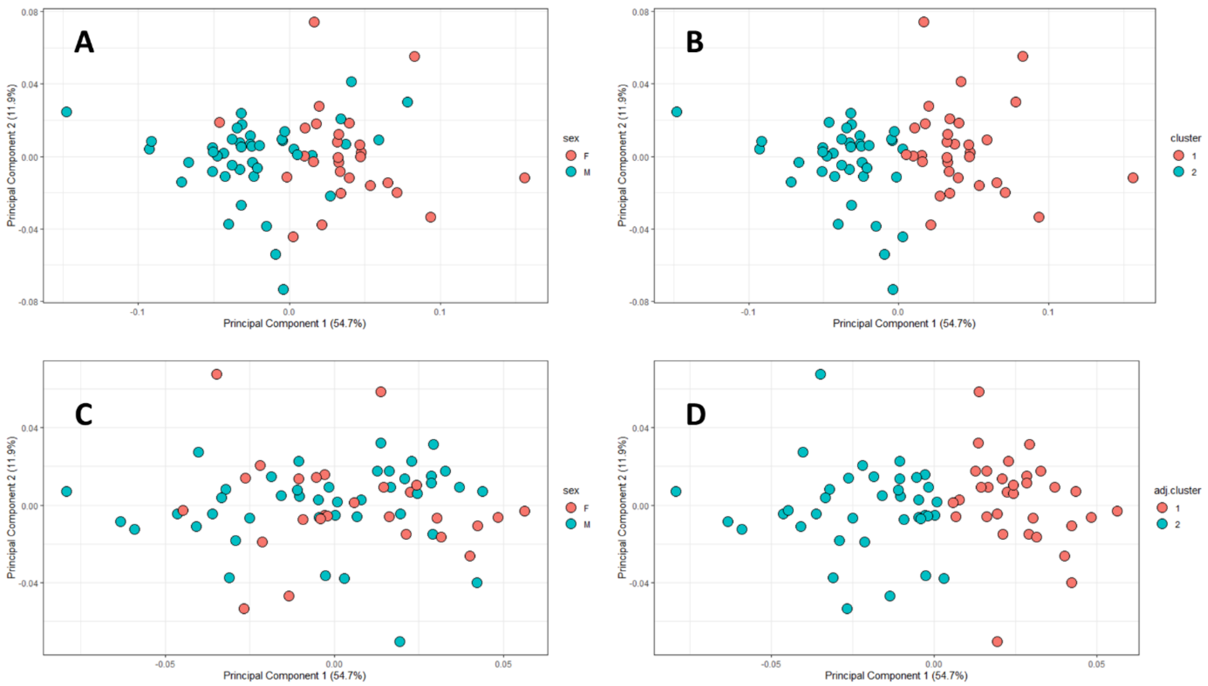


**Figure S5: Results of *K*-means cluster analysis of sex for n = 2 clusters, whole—skull shape data.** Shown are PCAs of **A:** Raw shape data, coloured by known sex; **B:** Raw shape data, coloured by cluster analysis estimate of sex; **C:** Allometry-corrected shape data, coloured by known sex; **D:** Allometry-corrected shape data, coloured by cluster analysis estimate of sex.

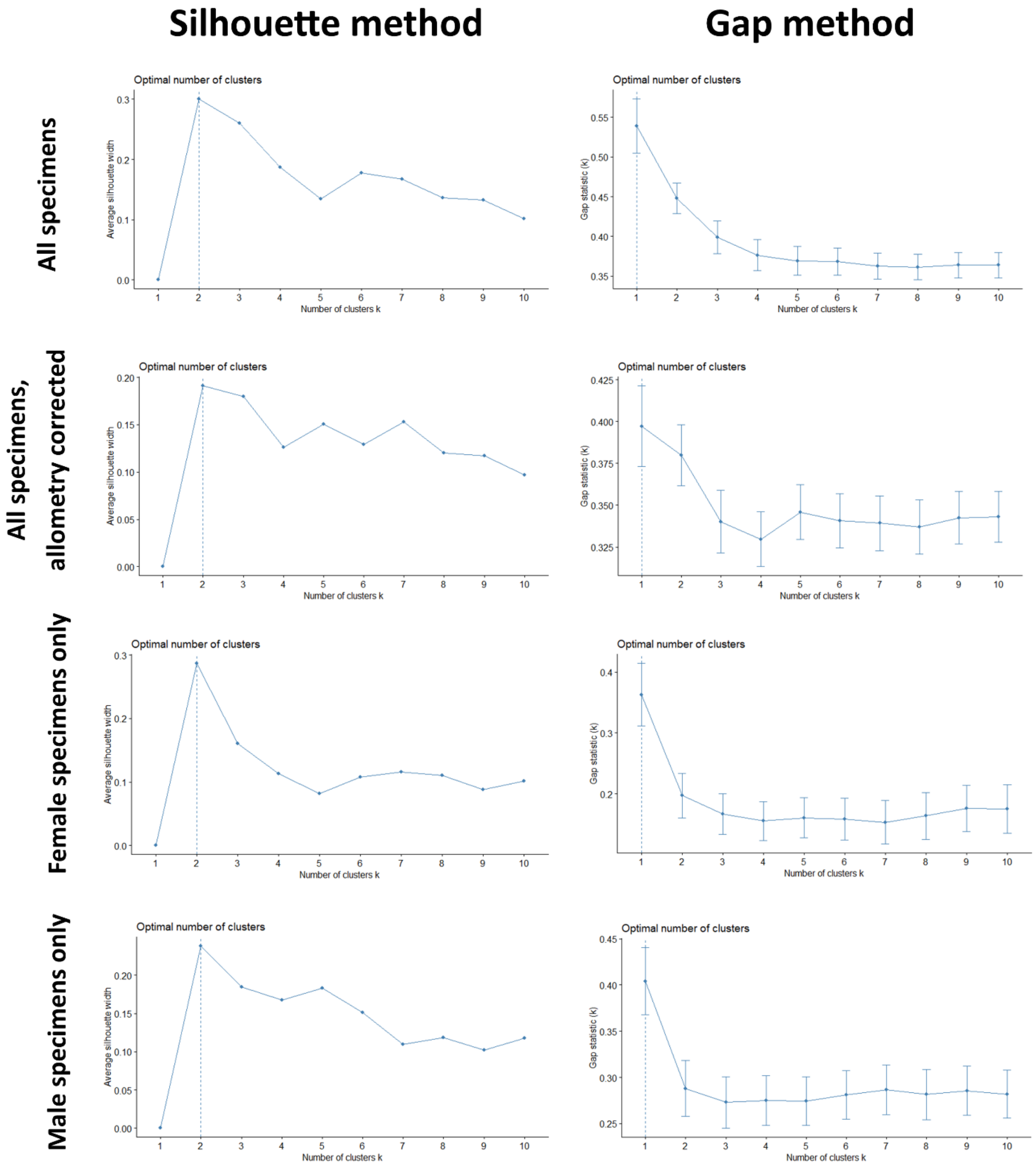


**Figure S6: Results of optimum cluster number analysis.** Left column shows results of 'silhouette' method for (top to bottom) all specimens (n = 70), allometry corrected specimens (n = 70), female specimens only (n = 27) male specimens only (n = 43). Right column shows results for 'gap' method for same subdivisions of dataset. Vertical dashed line in each plot represents calculated optimum cluster number.

**Table S9: Morphological variance of whole skull and individual modules, for all data and separate sexes.**

| | Raw data | | | Allometry corrected | | |
|---|---|---|---|---|---|---|
| | **All** | **Males** | **Females** | **All** | **Males** | **Females** |
| **Whole skull** | 5.10 | 4.22 | 3.72 | 3.15 | 3.15 | 2.82 |
| **Face (compare.CR)** | 1.95 | 1.62 | 1.60 | 1.43 | 1.28 | 1.42 |
| **Cranium (compare.CR)** | 1.88 | 1.89 | 1.42 | 1.46 | 1.53 | 1.19 |
| **Horn** | 14.61 | 11.66 | 10.29 | 8.24 | 8.40 | 7.33 |
| **Nasal** | 1.61 | 1.34 | 1.43 | 1.26 | 1.02 | 1.32 |
| **Premaxilla/maxilla** | 2.41 | 1.90 | 1.86 | 1.67 | 1.48 | 1.67 |
| **Frontal/parietal** | 1.62 | 1.52 | 1.33 | 1.31 | 1.30 | 1.15 |
| **Zygomatic** | 1.52 | 1.40 | 1.52 | 1.28 | 1.20 | 1.29 |
| **Lacrimal** | 1.01 | 0.97 | 0.94 | 0.90 | 0.88 | 0.84 |
| **Temporal** | 1.96 | 2.21 | 1.28 | 1.60 | 1.85 | 1.05 |
| **Occipital/sphenoid** | 2.27 | 2.30 | 1.65 | 1.64 | 1.72 | 1.34 |
| **Palatine** | 1.19 | 1.22 | 0.89 | 0.86 | 0.97 | 0.68 |
